# Screening, Identification and Growth Promotion Ability of Phosphate Solubilizing Bacteria from Soybean Rhizosphere under Maize-Soybean Intercropping Systems

**DOI:** 10.1101/2020.12.15.422997

**Authors:** Wenjing Wang, Clement Kyei Sarpong, Chun Song, Xiaofeng Zhang, Yuefeng Gan, Xiaochun Wang, Taiwen Yong, Xiaoli Chang, Yu Wang, Wenyu Yang

**Affiliations:** Institute of Ecological and Environmental Science, College of Environmental Science, Sichuan Agricultural University, Chengdu 611130, China; Sichuan Engineering Research Center for Crop Strip Intercropping System, Chengdu 611130, China; College of Landscape Architecture, Sichuan Agricultural University, Chengdu 611130, China

**Keywords:** intercropped soybean, phosphorus solubilizing bacteria, indole acetic acid, siderophore, growth promotion ability

## Abstract

The solubilization and mineralization of phosphorus by phosphate-solubilizing bacteria (PSB) is one of the most important bacterial physiological characteristics in the soil biogeochemical cycle. Through the isolation and screening of microorganisms in the rhizosphere soil of intercropped soybean in Ya’an, Renshou and Chongzhou, 9 PSBs with high phosphorus solubilizing ability were identified. It mainly belongs to *Bacillus* and *Pseudomonas*. The phosphate solubility of *Bacillus aryabhattai* B8W22 is as high as 388.62 µg·mL^-1^. The physiological and biochemical characteristics of each strain showed that it can secrete organic acids such as formic acid, acetic acid lactic acid and pyruvic acid. In addition, all strains can produce indole acetic acid and siderophores that promote plant growth. Seed germination experiments also showed that the phosphorus solubilizing bacteria isolated in this research have a certain ability to promote plant growth.

**IMPRTANCE:** *Bacillus aryabhattai* from rhizosphere soil of intercropped soybean has high phosphate-solubilizing ability, could produce indole acetic acid and siderophores that promote plant growth, and are of great significance in reducing the application of chemical phosphate fertilizers and promoting sustainable agricultural development.

## INTRODUCTION

Phosphorus is one of the most important nutrient elements for plant growth and development, and it plays a vital role in plant metabolism and various functions (Kochian, 2012). Phosphorus exists in various forms in the soil, but a large proportion exists in an insoluble form, due to fixation and precipitation by soil minerals such as calcium, aluminum, iron, and manganese (Pereira and Castro, 2014). It is estimated that the low availability of phosphorus affects more than 2 billion hectares of land worldwide (Shenoy and Kalagudi, 2005). Phosphorus deficiency leads to a significant decrease (5%-15%) in crop yields (Wissal et al., 2020). Plant suffering from phosphorus deficiency show symptoms of leaf redness and necrosis of old leaf tips. Phosphorus is usually the largest most limiting element for plant growth and development. To ensure sufficient and easily available form of phosphorus nutrient for optimal plant growth, phosphorus nutrients are usually supplemented to crop in the form of chemical fertilizers. However, intensive applications in recent years has/and is degrading ground and surface water bodies through soil erosion, surface runoff and groundwater leaching of residual phosphorus from agricultural fields (Bai et al., 2013). It is estimated that more than 15 million tons of phosphate fertilizers are used worldwide each year, and up to 80% of them are lost in an insoluble form (Gyaneshwar et al., 2002).

Under the current background of advocating ecological agriculture, excellent cultivation, and management measures are being adopted to increase the nutrient (P) absorption efficiency of crops by inoculating plant growth-promoting rhizobacteria (PGPR) (Billah et al., 2019). PGPR are beneficial microorganisms that live in the soil or epiphyte in the roots of plants, promote nutrient absorption and growth of plants, and inhibit harmful organisms (Bishnoi, 2015). PGPR represents a variety of soil bacteria, some of which can promote plant growth and development through nitrogen fixation, phosphorus solubilization, potassium solubilization and secretion of plant hormones such as indole acetic acid, etc., and can also enhance plants’ resistance to diseases, heavy metals, resistance to adversity such as salt and alkali stress (Mahanty et al., 2017).

It is estimated that under favorable conditions, the application of phosphate-solubilizing bacteria (PSB) can reduce the application of phosphate fertilizer by 50%, and will not significantly reduce crop production (Manzoor et al., 2016). PSB can synthesize low molecular weight organic acids, such as gluconic acid and citric acid. These organic acids have hydroxyl and carboxyl groups, which can chelate the cations bound to the phosphate, thereby converting insoluble inorganic phosphorus into its soluble form. Further, phosphate solubilizers synthesize different phosphatases to catalyze the hydrolysis of phosphates to promote the mineralization of organophosphates (Dastager and Damare, 2013).

Intercropping is a kind of traditional Chinese agriculture with a long history, can not only improve the index of multiple cropping of arable land (Yang et al., 2013), increase grain production and farmers’ income (Song et al., 2020), but also effectively alleviate the problems of low fertilizer use efficiency and negative environmental effect (Foley et al., 2011; Gebru, 2015; Yin et al., 2020). A large number of studies have shown that due to the complementary and mutually beneficial effects of crop species, intercropping between gramineous crops and leguminous crops can increase the utilization of nutrient resources and affect the biological characteristics of rhizosphere soil (Li et al., 2014). In terms of phosphorus nutrients, leguminous crops can promote the absorption of phosphorus by neighboring gramineous crops, change the phosphorus content of various forms of rhizosphere soil, and improve phosphorus utilization (Xue et al., 2016).

Maize-soybean relay strip intercropping is a major planting pattern in southwestern China (Yang et al., 2015). Soybean-based maize strip intercropping increases the phosphorus availability and APase activity in the soil thus improving soil fertility and production of the following crop (Du et al., 2018). In previous studies, the utilization rate of phosphate fertilizer in the maize-soybean intercropping system reached 84.6%, 20.4% higher than that of single-cropping maize (Song et al., 2020). Thus, the improvement in the agronomic performance of the applied P fertilizer under the maize-soybean intercropping system might have been due to the presence of efficient PSBs in the soybean rhizosphere that can release different types of organic acids and enzymes to promote the dissolution of insoluble phosphorus in the soil. It is therefore imperative that we screen and identify possible indigenous PSB under this cropping system that has the traits to facilitate phosphorus bioavailability and fertilizer use efficiency. In this study, we screened and identified PSB from soybean rhizosphere soil under maize-soybean intercropping systems in southwest China determined their phosphorus-solubilizing ability, physiological and biochemical characteristics, indole acetic acid secretion and siderophore production activity and further studied the plant growth promotion mechanism of PSB. The findings have important scientific significance and practical value in reducing the application of chemical phosphate fertilizers and promoting environmentally friendly agricultural production

## EXPERIMENTAL

### Methods

#### Soybean rhizosphere soil collection

Soybean rhizosphere soils were excavated from three experimental sites (Ya’an, Renshou, and Chongzhou) during the soybean vegetative growth phase under maize-soybean intercropping systems. For each sample, the roots of vigorously growing soybean were carefully dug. The bulk soils loosely bound on the roots were shaken off and discarded and the tightly-adhering soil to the soybean roots was moderately agitate in an Erlenmeyer flask filled with 100 mL sterile water. The roots were removed and the turbid liquid was used to directly isolate the PSB. Triplicate samples were used for all experiments

#### Medium and reagent formula

Pikovskaya’s agar (PVK) medium: glucose (10 g), Ca_3_(PO_4_)_2_ (5 g), (NH_4_)_2_SO_4_ (0.5 g), NaCl (0.2 g), KCl (0.2 g), MgSO_4_·7H_2_O (0.1 g), yeast extract (0.5 g), MnSO_4_ (0.002 g), FeSO_4_ (0.002 g), agar (18∼20 g), distilled water (1000 mL), pH 7.0∼7.2.

NBRIP liquid medium: glucose (10 g), Ca_3_(PO_4_)_2_ (5 g), MgCl_2_·6H_2_O (5 g), MgSO_4_·7H_2_O (0.25 g), KCl (0.2 g), (NH_4_)_2_SO_4_ (0.1 g), distilled water (1000 mL), pH 7.0.

Congo red liquid medium: Mannitol (10 g), K_2_HPO_4_·3H_2_O (0.5 g), MgSO_4_·7H_2_O (0.2 g), NaCl (0.1 g), yeast extract (1 g), NH_4_NO_3_ (1 g), L-tryptophan (0.1 g), 0.25% Congo Red (10 mL), distilled water (1000 mL), pH 7.0.

LB liquid medium: yeast extract (5 g), NaCl (10 g), Tryptone (10 g), distilled water (1000 mL), pH 7.0.

#### Isolation of PSBs

Each soil turbidity liquid was serially diluted. Aliquot of each dilution was spread on PVK medium and incubated at 28°C for 3-5 days. Colonies were selected on the basis of the development of phosphate-dissolving circles. Pick out single colonies with phosphate-dissolving circles and different morphologies, and perform plate streaking and purification. Once purified, each isolate was stored at −80°C in the same medium with 20% (v/v) glycerol.

#### Determination of phosphate-solubilizing ability

Inoculate the strains from the preliminary screening on PVK medium. Place the plate upside down in a constant temperature and humidity incubator at 28°C for 3-5 days, measure its phosphate-dissolving circle diameter (D) and colony diameter (d), and calculate the ratio (D/d). Select strains with larger ratio (D/d) for quantitative determination to determine their phosphate-solubilizing ability. The strains were inoculated into the NBRIP liquid medium at an inoculum of 1%. The suspension was incubated on a shaker at 180 rpm and 28°C for 5 days and then centrifuged at 8000 rpm, 4°C for 10 min. The water-soluble phosphorus content in the supernatant was determined by the molybdenum blue method (J. and P, 1962). Autoclaved un-inoculated medium was used as the control treatment. Use a pH meter to measure the change in pH of the suspension.

#### Strain identification

16S rDNA sequencing and phylogenetic analysis were performed for taxonomical assignment. The target strain was activated in LB liquid medium, and the bacterial genomic DNA extraction kit (Sangon Biotech Chengdu Co., Ltd.) was used to extract the genomic DNA of the strain. The rDNA fragments of 16s rDNA were amplified with the universal primer 27F (5’-AGA GTT TGA TCC TGG CTC AG-3’) and 1492R (5’-GGT TAC CTT GTT ACG ACTT-3’) (GUPTA et al., 1994). Polymerase chain reaction (PCR) amplification which consists of a 50 µL system, including template DNA 2 µL, forward primer 2 µL, reverse primer 2 µL, 2×TaqPCR premixed enzyme 25 µL, and DdH_2_O 19 µL was performed as follows: 95°C for 5 min in initial denaturation, 30 cycles of 95°C for 30 s, 57°C for 30 s, 72°C for a 1 min denaturation annealing and extension, and 72°C for a 5 min final extension of the amplified DNA. The amplified products from three technical replicates were quantified with 1% agarose gel electrophoresis and mixed in one tube and were sent to Chengdu Youkang Jianxing Biotechnology Co., Ltd. for sequencing. The sequences were compared against the GenBank database using the NCBI BLAST program. The phylogenetic tree was constructed using MEGA 7.0 software. The sequences were deposited into GenBank and the accession numbers were obtained.

#### Analysis of indole acetic acid production

Indole acetic acid (IAA) production of PSBs was determined according to the method of Gordon and Weber. (1951) with some modifications. The test strains were inoculated into L-tryptophan-containing LB liquid medium (addition of 1 L LB + 0.1 g L-tryptophan). The suspension was incubated on a shaker at 180 rmp at 28°C for 2 days. The broth was centrifuged at 8000 rpm for 10 min. The supernatant was reserved and 0.1 mL was mixed with o.1 mL of Salkowski’s reagent (4.5 g FeCl_3_ and 587.4 mL 98%H_2_SO_4_, solution) and kept in the dark for 15 min. Use an enzyme-labeled instrument to measure the absorbance at a wavelength of 530 nm. Calculate the content of IAA per unit volume of fermentation broth according to the standard curve.

#### Analysis of siderophore production activity

All the test strains were respectively inoculated into MSA liquid medium, shaking culture at 28°C, 180 rmp incubator for 2 days. The fermentation broth was centrifuged at 8000 rmp for 10 min. The production of siderophore by the isolate was quantitatively evaluated using Chrome Azurol sulphonate (CAS) assay as described by Shin et al. (2001). CAS solution consisting of 60.5 mg of CAS was diffused in 50 ml of deionized water to which 10 ml of FeCl_3_·6H_2_O the solution was added. 72.9 mg HDTMA (Hexa-decyl Trimethyl Ammonium bromide) dissolved in 40 ml of deionized water was added to CAS to make the volume to 100 ml. From the prepared CAS solution, 4 ml was taken and mixed thoroughly with 4 ml of the culture supernatant and was incubated for 1 hour. The absorbance of each sample was measured at a wavelength of 630 nm. Siderophore production by bacterial strains was evaluated as the percentage of siderophore units (SUs=[(Ar-As)/Ar] ×100) where Ar = absorbance of reference (CAS reagent); As = absorbance of the sample (Payne, 1993).

#### Analysis of physiological and biochemical characteristics of strains

The physiological and biochemical characteristics of each strain were analyzed by the catalase test, starch hydrolysis test, indole test, methyl red test, V-P test, hydrogen sulfide test, urea test and gelatin liquefaction test.

#### Analysis of seed germination percentage

To evaluate the percent germination, homogenous sterilized deionized water-washed maize seeds were soaked in fermented broth for 30 min. The fermented broth was prepared by culturing the bacteria strains in 100 mL of LB medium at 28°C for 36 h, and then cells were harvested by centrifugation at 5000 rpm at 4°C for 10min and adjusted to 1 × 10^8^ cells/mL in sterile deionized water. The seeds were propagated in a petri dish covered with a layer of moist filter paper and incubated at 25°C. 2mL of sterile water was added to each treatment on a daily basis.

#### Statistical Analysis

The statistical data analysis was carried out by IBM^®^ SPSS^®^ Statistics V. 26 software. One-way ANOVA (analysis of variance) was used, followed by LSD(L) and Waller-Duncan(W) post hoc test to determine the significant difference among means of the treatment at 0.05 significance level.

## RESULTS AND DISCUSSION

### Isolation and Phosphate-solubilizing Ability of PSBs

Initially, 44 strains with halo zones in PVK agar medium were isolated as the positive microbes, demonstrating phosphate solubilize ability. Among them, 14 strains were isolated from Ya’an soil samples, 14 strains were isolated from Chongzhou soil samples, and 16 strains were isolated from Renshou soil samples. According to the analysis of the diameters of the phosphate-dissolving circle (Table 1) and the content of available phosphorus (Fig.1), a total of 9 strains with better phosphate-solubilizing ability were selected for the follow-up study. The results showed that all the 9 strains could solubilize Ca_3_(PO_4_)_2_ in large quantities. The ratio of the diameter phosphate-dissolving circle to the diameter of colony (solubility index) of Y5, C8, C9, and C10 was significantly higher than other strains. The soluble phosphorus content in the supernatant of strains Y7, Y9, R7, and C9 were relatively high with 380.96 µg·mL^-1^, 388.62 µg·mL^-1^, 374.48 µg·mL^-1^, 381.30 µg·mL^-1^, respectively.

**Table 1.** The diameters of the colonies and phosphate-dissolving circle of PSBs and their ratios.

| Strains | The diameters of the colonies (d, mm) | The diameters of the phosphate-dissolving circle (D, mm) | Solubility index (D/d) |
| --- | --- | --- | --- |
| Y5 | 2.50±0.31d | 10.81±0.64c | 4.38±0.66a |
| Y7 | 3.14±0.34c | 9.69±1.06d | 3.10±0.33f |
| Y9 | 3.05±0.36c | 10.27±0.51cd | 3.39±0.33df |
| R1 | 3.72±0.21ab | 13.15±0.69ab | 3.54±0.21cdf |
| R7 | 4.11±0.13a | 13.65±0.67a | 3.33±0.17f |
| R9 | 3.47±0.49bc | 13.11±0.53ab | 3.83±0.41bcd |
| C8 | 3.45±0.25bc | 13.67±0.55a | 3.97±0.19abc |
| C9 | 3.11±0.16c | 13.24±0.66ab | 4.27±0.22ab |
| C10 | 3.27±0.26c | 12.83±0.53ab | 3.95±0.31abc |

Y5, Y7 and Y9 were isolated from Ya'an soil samples; R1, R7 and R9 were isolated from Renshou soil samples; C8,

**Fig.1.**
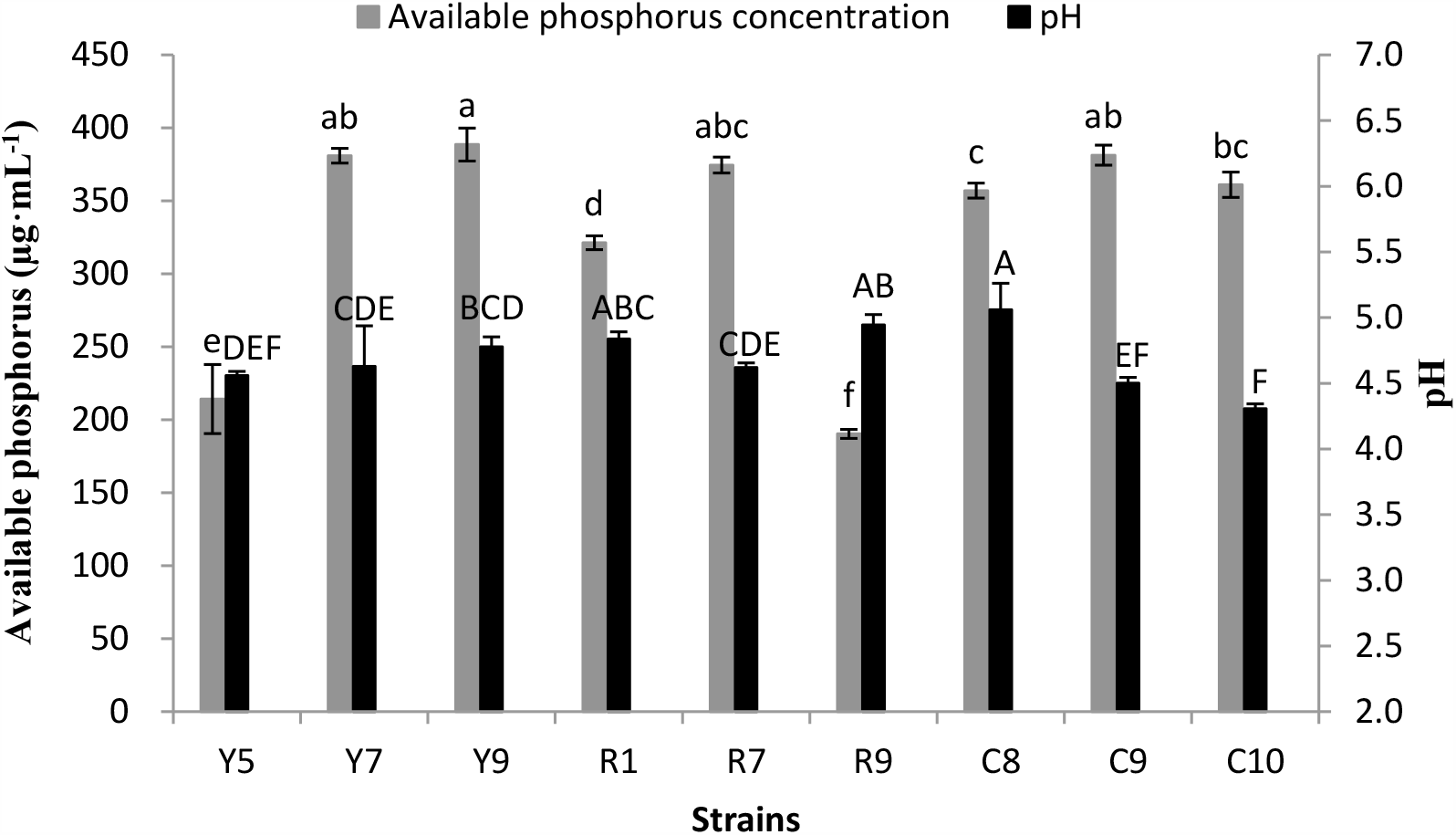
The content of available phosphorus and pH in supernatant of strains cultured in NBRIP liquid medium for 5 d. Error bars represent the standard deviation of three replicates. Mean values labeled with the same letter were not significantly different at *p* < 0.05.

### Morphological Characteristics of PSBs

The morphological characteristics of PSBs are shown in Table 2. The shape and color of each strain are similar, only the cell shape of the bacteria is slightly different. Using Gram staining method, observe the morphology of bacteria with an optical microscope. The results showed that besides strain R1 is a gram-positive bacterium, all the other strains are gram-negative bacteria.

**Table 2.**
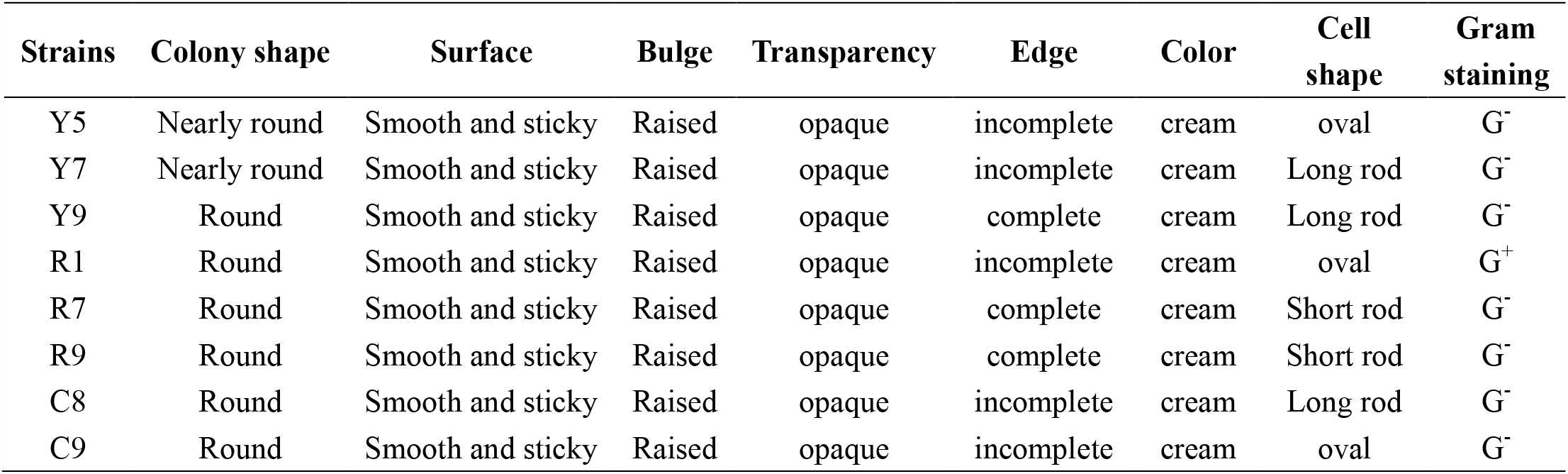
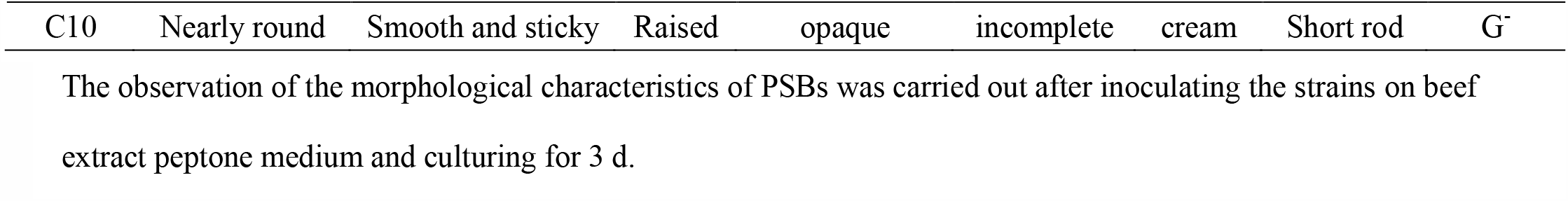
Morphological characteristics of PSBs.

### Molecular Identification of PSBs

Molecular identification was conducted with MEGA 7.0 software using a neighbor-joining method, and sequence analysis was conducted with DNAstar 7.1 software. The phylogenetic tree is shown in Fig.2. Strain Y5 was identified as *Novosphingobium resinovorum* NCIMB 8767 (NR044045.1), strain Y7 was identified as *Kosakonia sacchari* SP1 (NR118333.1), strain Y9 was identified as *Bacillus aryabhattai* B8W22 (NR115953.1), strain R1 was identified as *Bacillus thaonhiensis* NHI-38 (NR125615.1), Strain R7 was identified as *Bacillus simplex* LMG 11160 (NR114919.1), strain R9 was identified as *Bacillus subtilis* JCM 1465 (NR113265.1), strain C8 was identified as *Enterobacter cloacae* DSM 300549 (NR117679.1), strain C9 was identified as *Ralstonia pickettii* NBRC 102503 (NR114126.1), and strain C10 was identified as *Pseudomonas plecoglossicida* NBRC 103162 (NR114226.1).

**Fig.2.**
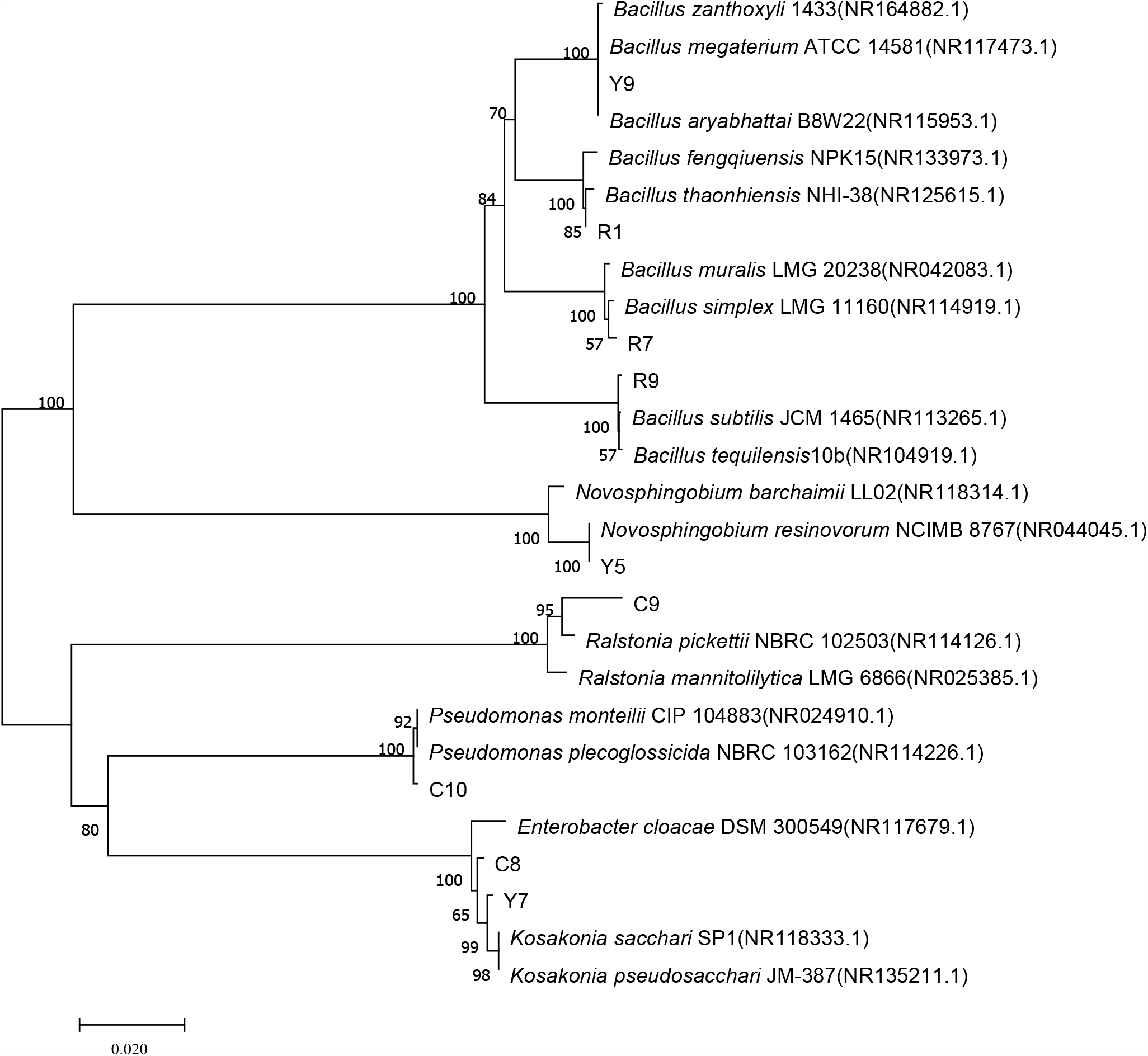
Phylogenetic tree of established based on 16S rDNA sequences of 10 phosphate-solubilizing bacteria.

### Physiological and Biochemical Characteristics of PSBs

The main physiological and biochemical characteristics of each strain are shown in Table 3. The results showed that some bacteria can secrete some extracellular enzymes such as catalase, hydrolase, tryptophanase, urease, and protease. These enzymes metabolize substances in vitro to meet the self-growth needs of strains. The methyl red test and V-P test of all strains were positive, indicating that all strains can use glucose to produce organic acids, such as formic acid, acetic acid, lactic acid, and pyruvic acid. Inorganic phosphorus solubilization occurs as a result of the action of low molecular weight organic acids such as gluconic and citric acids which are synthesized by different soil bacteria (Keshav et al., 2013). These organic acids have hydroxyl and carboxyl groups, which can chelate the cations that bind to phosphates, resulting in the conversion of insoluble phosphorus to its soluble form (Mahanty et al., 2017).

**Table 3.** Physiological and biochemical characteristics of selected strains

| Strains | Catalase test | Starch hydrolysis test | Indole test | Methyl red test | V-P test | Hydrogen sulfide test | Urea test | Gelatin liquefaction test |
| --- | --- | --- | --- | --- | --- | --- | --- | --- |
| Y5 | + | - | + | + | + | + | - | - |
| Y7 | + | - | + | + | + | + | + | - |
| Y9 | + | + | + | + | + | + | + | - |
| R1 | + | - | + | + | + | + | - | - |
| R7 | + | + | + | + | + | + | + | - |
| R9 | + | + | + | + | + | + | - | - |
| C8 | + | - | + | + | + | + | + | - |
| C9 | - | - | + | + | + | + | + | - |
| C10 | + | - | + | + | + | + | + | - |

"+" indicates a positive reaction, and "-" indicates a negative reaction. Catalase test, starch hydrolysis test, indole test, urea test and gelatin liquefaction test "+":indicates that the strain can secrete catalase, hydrolase, tryptophanase, urease and protease; methyl red test "+": indicates that bacteria can metabolize sugar to produce organic acids, such as formic acid, acetic acid, lactic acid, etc.; V-P test "+": indicates that bacteria can use glucose to produce non-acidic or neutral end products, such as pyruvic acid. Hydrogen sulfide test "+": indicates that the strain can decompose sulfur-containing organic matter such as cystine, cysteine and methionine.

### Indole Acetic Acid Production Ability of PSBs

The bacteria strains were examined for their acetic acid (IAA) production ability. IAA is a ubiquitous endogenous auxin in plants, mainly produced by the plant’s own meristem. Studies have shown that IAA can promote the growth and development of crops and the absorption and utilization of nutrients by promoting root hair growth and root elongation (Abel and Theologis, 2010; Suliasih and Widawati, 2020). As shown in Fig.3, all strains secreted IAA. Strain Y9, C9, and C10 secreted high quantities of IAA and Y9 recorded the highest (26.17 μg·mL^-1^) and it was significantly higher than both C9 and C10 which were statistically similar.

**Fig.3.**
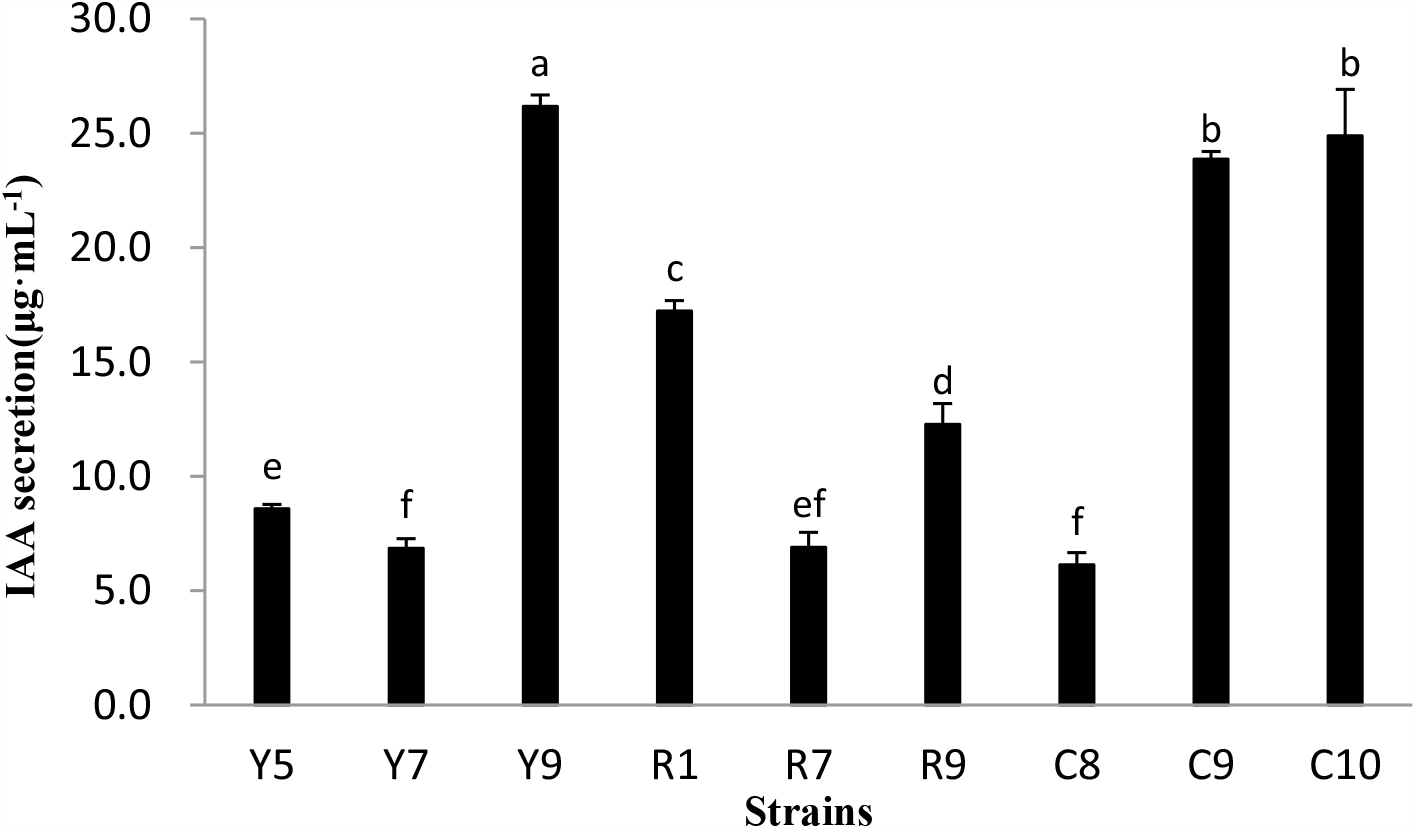
Determination of IAA secretion of different strains. Error bars represent the standard deviation of three replicates. Mean values labeled with the same letter were not significantly different at *p* < 0.05.

### Siderophore Production Activity of PSBs

Endophytic bacteria that produce siderophores can reduce the competitiveness of pathogens by providing available iron to host plants or reducing the available iron of pathogens in the environment, thereby promoting plant growth and development (Loper and Henkels, 1999). Siderophore is a low molecular weight substance that can specifically bind Fe^3+^ and provide microbial cells (Buyer et al., 1993; Ma et al., 2012). Iron is an essential element for the growth of microorganisms. In an iron-deficient environment, microorganisms secrete siderophores to chelate iron ions to meet their own growth needs (Wang et al., 2013). The siderophore production activity of each test strain was determined by Chrome Azurol S (CAS) method, and the result is shown in Fig.4. All strains have the ability to synthesize siderophores, among which strains C8 have the highest siderophore production activity, up to 64.5 units.

**Fig.4.**
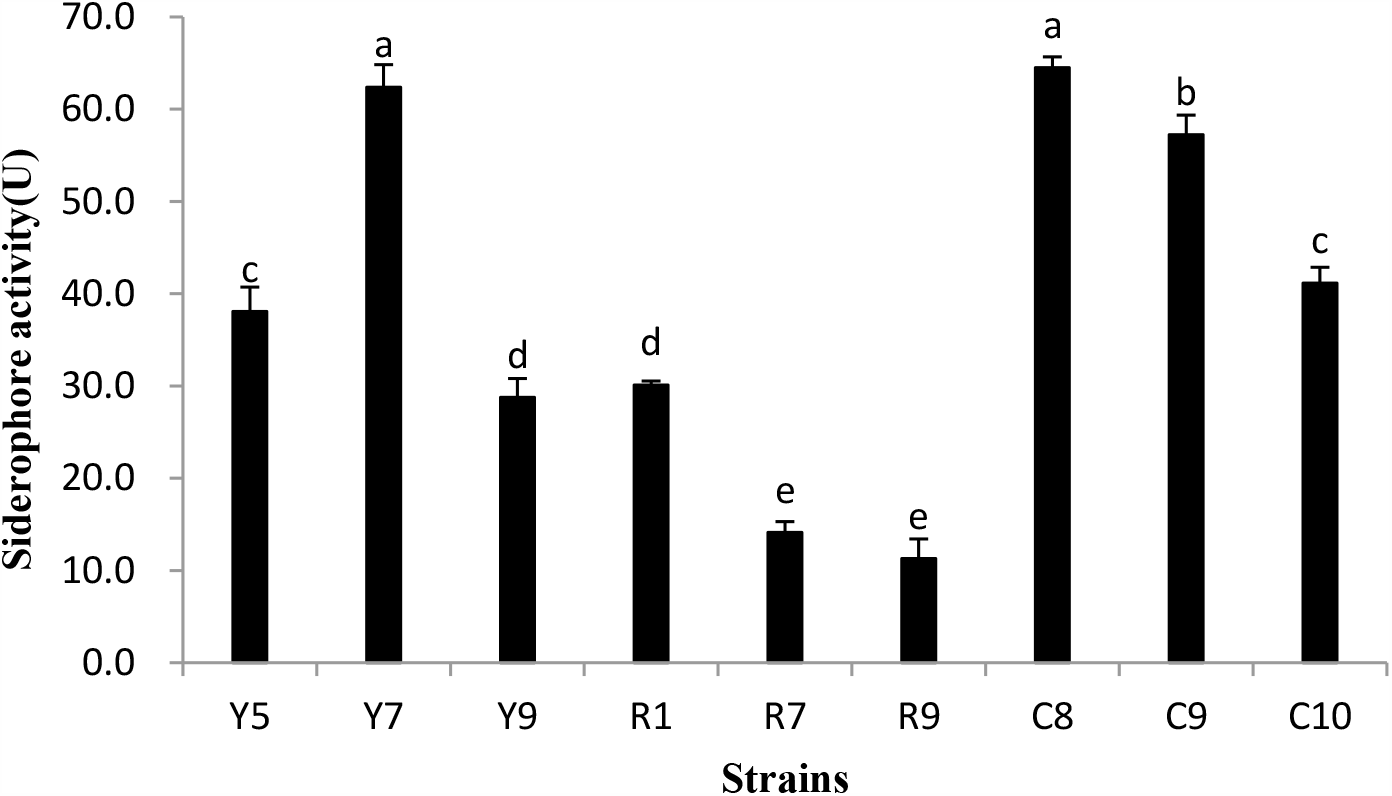
Determination of siderophore production activity of different strains. Error bars represent the standard deviation of three replicates. Mean values labeled with the same letter were not significantly different at *p* < 0.05.

### Analysis of Seed Germination Percentage

After maize seeds were cultivated in a light incubator for 4 days, the germination percentage and rooting number were counted. As shown in Fig.5, each strain can effectively increase the germination percentage of seeds. The stronger growth-promoting effects are strains Y9, R1, R9, C9 and C10. The germination rate of CK was 63.3%, while the germination rate of the experimental group treated with the bacterial solution was above 70%. There was no significant difference in the number of roots of maize seeds inoculated with each strain. The number of roots of seeds treated with sterile water was 2.86, and the number of roots of seeds inoculated with strains was 2.67-3.71.

**Fig.5.**
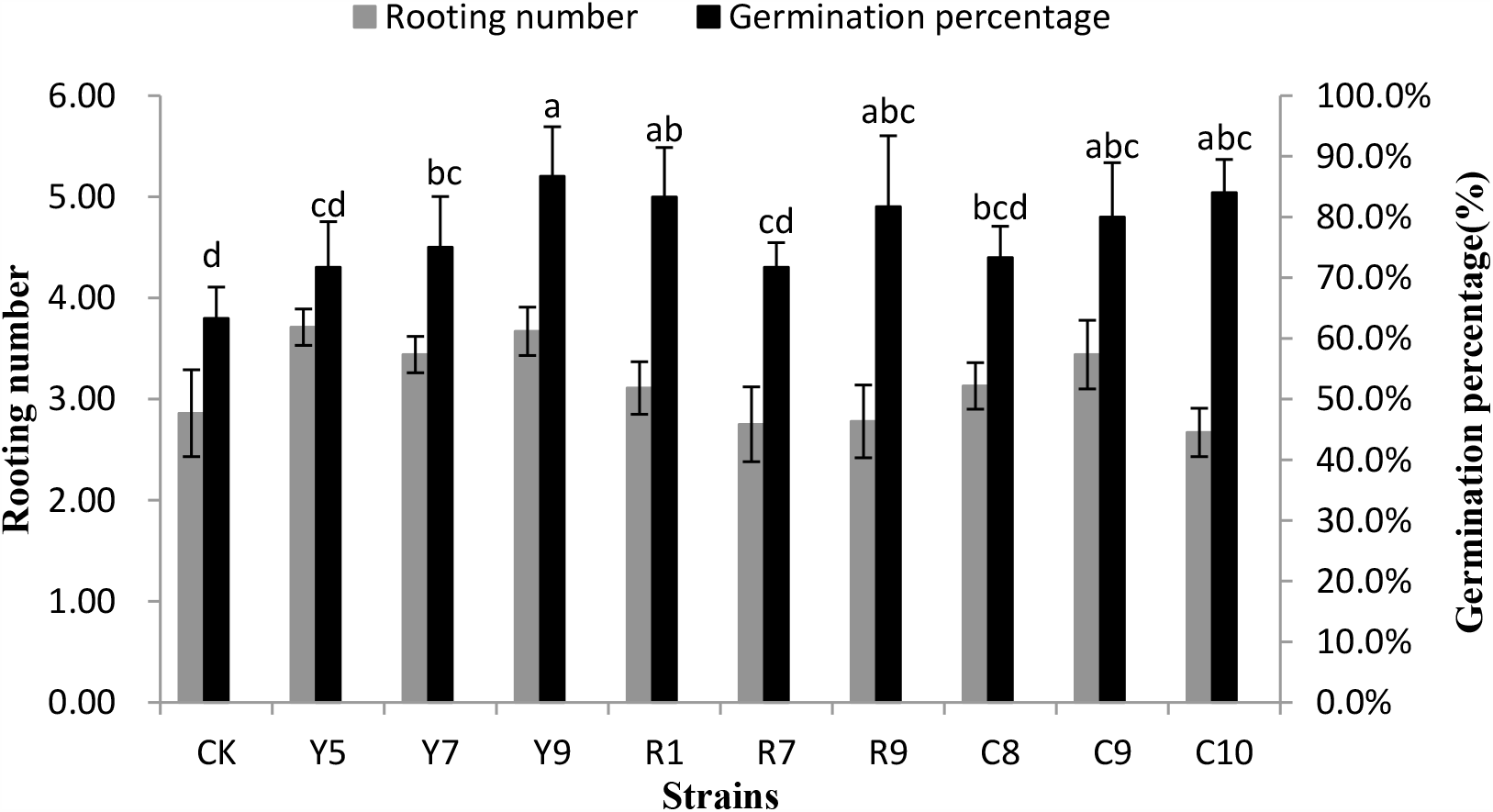
Germination percentage and rooting number of maize seed inoculated with different strains. CK: The seeds were soaked in sterile water as a blank control. Rooting number: The average number of roots per seed. Germination percentage: Refers to the percentage of germinated seeds to test seeds after 4 d of cultivation. Error bars represent the standard deviation of five replicates and each replicate consists of ten maize seeds per plate. Mean values labeled with the same letter were not significantly different at *p* < 0.05.

## Discussion

Soil microorganisms are an important part of the agroecosystem and play an important role in the recycling of soil nutrients, degradation of pesticides, and maintenance of soil health, sustainability, and productivity (Buyer et al., 2010). A plethora of microorganisms with phosphorus solubilizing function have been isolated from different soils including calcareous rhizosphere soil, acid sulfate soil, and corn, rice, and wheat rhizosphere (Ali et al., 2014; Liu et al., 2015; Manzoor et al., 2016). The population structure of phosphorus solubilizing microorganisms existing in the soil is mainly composed of phosphorus solubilizing bacteria, phosphorus solubilizing fungi and phosphorus solubilizing actinomycetes. The predominant PSBs that have been reported to demonstrating strong phosphate solubilizing abilities are *Bacillus* and *Pseudomonas* (Chung et al., 2005; Mander et al., 2012; Tilak et al., 2005; Zeng et al., 2016). Inoculating PSB strains induce the availability of poorly soluble phosphorus in the soil and increase the phosphorus absorption rate by crops. Hence facilitating the recycling of poorly soluble phosphorus in the soil, and can minimize the application of phosphate fertilizer (Billah et al., 2019).

PSB belonging to different genera can promote the absorption and growth of different crops of insoluble phosphorus (Vessey, 2003). However, most of the the research focused on maize, wheat, rice, and other gramineous plants. There are few studies on the use of legumes to verify the phosphorus solubilizing effect and the mechanism of phosphorus solubilization. In three different field studies on wheat (Abaid-Ullah et al., 2015; Afzal et al., 2005; Chen et al., 2006), it was observed that the application of phosphorus solubilizing bacteria (*Pseudomonas* and *Bacillus*) can increase plant nutrient accumulation, dry weight, and grain yield. In a greenhouse experiment of wheat (Selvakumar et al., 2009), the application of phosphorus solubilizing bacteria increased seed germination rate and plant biomass. In another study, (greenhouse and field experiments) on corn (Gholami et al., 2009; LEGGETT et al., 2015; Pandey et al., 2007; Yazdani et al., 2009), the application of phosphorus solubilizing bacteria (*Azobacterium, Pseudomonas, Serratia, Bacillus*, etc.) can increase the chlorophyll content, biomass, and yield of plants. Further, the application of phosphate solubilizing agents (*Serratia* and *Spirulina orphanata*) significantly increased the grain yield, root length, leaf length, and plant dry weight of rice (Araujo et al., 2013). In this study, new isolates (*Kosakonia, Enterobacter, Ralstonia*) demonstrated a strong capability to solubilize insoluble phosphate. A number of studies comparing continuous monocropping and intercropping systems shown that intercropping planting patterns improved biodiversity, land-use efficiency, and nutrient use efficiency (Cong et al., 2015; Wang et al., 2014; Zhu et al., 2000). This beneficial effect might be due to the complementary and mutually beneficial effects of crop species. Leguminous crops can promote the phosphorus absorption of adjacent gramineous crops by modifying the phosphorus content in the rhizosphere soil, and improve phosphorus utilization.

For example, intercropped peanut significantly increased phosphorus uptake compared with monocropped peanut in a corn-peanut intercropping system (Dessougi et al., 2003); In another study (wheat-chickpea intercropping system), the rhizosphere of chickpea secreted more acid phosphatase and hydrolyzed organic phosphorus into soluble inorganic phosphorus in the soil, thus promoting the absorption of phosphorus by wheat (Li et al., 2003; Li et al., 2004). Further, in a maize-pigeon pea intercropping system, soil P availability and organic phosphorus storage were enhanced through biochemical and physical rhizosphere interactions (Garland et al., 2017). The maize and soybean strip intercropping technology widely promoted in southwestern China can not only obtain higher crop yields, but also improve soil nitrogen and phosphorus utilization and microbial diversity (Liu et al., 2017; Liu et al., 2018).

Ben Zineb et al. (2019) obtained of the 8 strains that were from seven different soils with the phosphate-solubilizing ability ranged from 107 µg·mL^-1^ to 715 µg·mL^-1^ with TCP as phosphorus source and from 9.2 µg·mL^-1^ to 605 µg·mL^-1^ with TRP. Elhaissoufi et al. (2020) isolated 42 PSBs from the rhizosphere soils of several crops (wheat, barley, maize, oat, faba beans, peas, etc.), which showed a low phosphate-solubilizing ability ranging from 40 µg·mL^-1^ to 113 µg·mL^-1^. In our study, a total of 44 PSBs was isolated from the soybean rhizosphere soil samples under maize-soybean intercropping systems at three experimental bases of Yaan, Renshou, and Chongzhou. Among them, 9 strains had higher phosphate-solubilizing ability, ranging from 190 to 390 µg·mL^-1^.

The phosphate-solubilizing ability of PSB is affected by many environmental factors, such as soil temperature, salinity, pH, and dissolved oxygen (Wei et al., 2016; Zeng et al., 2016). The phosphorus solubilizing mechanism of PSB in the rhizosphere of crops is different in different regions, different soils and different planting patterns. Generally, the main reason for the dissolution of poorly soluble phosphate is that phosphorus solubilizing bacteria release acidic substances to lower the soil pH. The pH of PSBs isolated from the rhizosphere of local fragrant rice in the Bada River Valley, Sulawesi, Indonesia, was between 4.27 and 5.67 after 7 days of cultivation in liquid Pikovskaya media (Sudewi et al., 2020). The pH of 76 PSBs isolated from agricultural fields (Hailun, Heilongjiang, China and Yingtan, Jiangxi, China) were mostly between 4.37 and 8.34 after 3 days of cultivation in liquid modified PVK (without agar and indicator) (Zheng et al., 2018). In this study, the pH of all strains was between 4.31 and 5.06 after 5 days of cultivation in NBRIP medium,at a low level compared to other studies. The phosphorus solubilization mechanism of PSB is related to the decrease of soil pH through protons discharged from NH_4_^+^ assimilation, which lowers the pH value and causes the dissolution of phosphate (Chen et al., 2006). Further, the release CO_2_ through respiration can lower soil pH which causes the dissolution of phosphate (Ismail et al., 2016). Again, organic acid is synthesized through metabolic activities can directly dissolve insoluble phosphate and chelate mineral-phosphate bonds (Jones and Oburger, 2011; Zhang et al., 2014).

Furthermore, the multiple beneficial effects of PSB are widely regarded as the key factors to promote plant growth and increase the availability of soil phosphorus. Plant hormones or phytohormones play a crucial role in plant growth and development (Peleg and Blumwald, 2011). It is reported that 80% of microorganisms which have been isolated from the rhizosphere of various crops possess the ability to synthesize and releasing auxins as secondary metabolites (Ahemad and Khan, 2011). Most of the bacteria with significant growth-promoting effects produce IAA in the range of 1.47-32.8μg·mL^-1^ (AHMAD et al., 2005). The amount of IAA secreted by strain Y9 in this study is as high as 26.17 μg·mL^-1^. Besides IAA, iron is one of the important nutrients in almost all life forms. All plants, microorganisms, and animals need iron (Rajkumar et al., 2010). All strains can produce siderophore in this study, the siderophore production activity of the 9 PSBs ranged from 11.29 to 64.50 units. The siderophore production activity of ten bacterial isolates from wheat (rhizosphere soil and root endosphere) ranged from 31.71 to 53.71 units (Sood et al., 2018), which is similar to this work.

The IAA secreted by the strain has a certain effect on the germination and growth of plant roots. Therefore, the use of specific strains of fermentation broth to treat seeds and study its effect on seed germination is a factor in evaluating the growth-promoting effect of bacteria. In the seed germination experiment of this study, it has been preliminarily verified that PSBs can promote the growth of maize seeds to some extent. The results showed that after treatment with strains Y9, C10, and R1, the germination rate of maize seeds increased by 23.4%, 20.7%, and 20.0%, respectively, compared with CK.

## CONCLUSIONS

A total of 9 strains of bacteria with high phosphate-solubilizing ability were identified. Y9, R1, R7, and R9 belong to *Bacillus*, and Y5, Y7, C8, C9, and C10 belong to *Novosphingobium, Kosakonia, Enterobacter, Ralstonia*, and *Pseudomonas*, respectively.

The strains could effectively utilize Ca_3_(PO_4_)_2_, and the water-soluble phosphorus content of strains Y7, Y9 and C9 were as high as 380.96 µg·mL^-1^, 388.62 µg·mL^-1^ and 381.30 µg·mL^-1^, respectively.

The strains were able to reduce pH to about 4. The physiological and biochemical characteristics of the strains showed that the strains can produce organic acids such as formic acid, acetic acid, lactic acid and pyruvic acid.

All the strains secreted IAA and produce siderophores. The content of IAA secreted by strain Y9 was as high as 26.17 μg·mL^-1^. Strains C8 have the highest siderophore activity, up to 64.5 units.

Seed germination experiments show that PSB can promote plant growth. Compared with the control group, the seeds treated with PSB have a higher germination rate and rooting number.

## ACKNOWLEDGMENTS

The work was funded by National Natural Science Foundation of China (31771726). Any opinions, finding, and conclusions or recommendations expressed in this material are those of authors and do not necessarily reflect the views of National Science Foundation of China.

